# *CREBBP* and *WDR 24* Affects Quantitative Variation in Red Colouration in the Chicken

**DOI:** 10.1101/281840

**Authors:** J. Fogelholm, R. Henriksen, A. Höglund, N. Huq, M. Johnsson, R. Lenz, P. Jensen, D. Wright

## Abstract

Plumage colouration in birds is important for a plethora of reasons, ranging from camouflage, sexual signaling, and species recognition. The genes underlying colour variation have been vital in understanding how genes can affect a phenotype. Multiple genes have been identified that affect plumage variation, but research has principally focused on major-effect genes (such as those causing albinism, barring, and the like), rather than the smaller effect modifier loci that more subtly influence colour. By utilizing a domestic x wild advanced intercross with a combination of classical QTL mapping of red colouration as a quantitative trait and a targeted genetical genomics approach, we have identified five separate candidate genes (*CREBBP, WDR24, ARL8A, PHLDA3, LAD1*) that putatively influence quantitative variation in red colouration in chickens. Such small effect loci are potentially far more prevalent in wild populations, and can therefore potentially be highly relevant to colour evolution.

## Introduction

The presence of feathers is one of the most defining characteristics of birds. Feathers are not only specialized for flight but also to relay a plethora of visual signals to both friends and foes alike through specific visual patterns, such as camouflage (GLUCKMAN AND CARDOSO 2010), sexual signaling and displays which can influence sexual selection (WILKINS *et al.* 2016) and species recognition (SEDDON *et al.* 2013). One of the most striking aspects of the pattern is the colouration of the plumage, which can range from pure black or white to metallic greens and purples as well as oranges and reds. The colour of the feathers can be sourced from the environment,(LOPES *et al.* 2016) or synthesized internally with genetic regulation. The most common colours are produced by the melanins. These pigments come in two forms: eumelanin, which produces blacks and browns (GALVÁN AND SOLANO 2016) and pheomelanin, which produces reds, oranges and yellows that are similar in appearance to the carotenoid pigments(GALVÁN AND SOLANO 2016)

Domesticated animals are a classical example of extreme variability in pigmentation within species. A reduction in pigmentation and colouration complexity is usually considered as one of the key components of the domestication syndrome (WILKINS *et al.* 2014), however artificial selection has also led to novel colour phenotypes in domesticated animals (LUDWIG *et al.* 2009; LINDERHOLM AND LARSON 2013; WRIGHT 2015). In the case of coat colouration, domestic animals are generally speaking lighter versions of their wild ancestors. This is for instance the case with chickens, where the wild ancestor the Red Junglefowl has a wide range of plumage colours ranging from dark red to light orange. Several of the domesticated chicken breeds such as the white leghorn layer and commercial broiler breeds provide a stark contrast to the colourful Red Junglefowl as they are completely white and display no sexual dichromatism.

There are several genes implicated in plumage colour variation. We can subdivide colour genes into three distinct classes depending on their mechanism. Intra-individual patterning genes determine the patterns we can observe across an individual. Intra-feather patterning genes determine the pattern within each feather (e.g barred feathers(SCHWOCHOW THALMANN *et al.* 2017)). Finally we have the colour determinants. These are genes that set the colour for each position within the observed pattern. This can be either in a single feather or across an individual. As an example of the independence of these types of colour loci, a recent study in a hemimetabolous insect (LIU *et al.* 2016), concluded that the genes influencing the patterning of the colours did not alter the intensity of the pigmentation. This suggests that coat colour patterns are achieved through a combination of patterning genes coupled to the genes that determine the intensity of each colour.

Examples of causative genes for pigmentation include well studied colour determining genes such as *MC1R* (involved in the melanin pathway where its activation leads to the production of black eumelanin (HEARING 2011)) and *MITF,* which activates the enzyme Tyrosinase (YASUMOTO *et al.* 1994), a rate limiting step in the melanin synthesis pathway. These variants can all be considered of major effect, in that they cause albinism, black plumage, etc in a binary fashion (though often a variety of epistatic interactions also occur). However to the best of our knowledge, smaller effect loci that have less extreme modifier effects, in essence quantitative trait loci for plumage colour, have yet to be identified. In wild individuals it may be of greater importance to more subtly modify coat colouration, which is why many of the genes with more extreme phenotypic effects are identified in laboratory or domestic populations.

To identify genes that regulate quantitative variation in the intensity of red colouration, we utilized an advanced intercross between wild Red Junglefowl and domestic White Leghorn chickens. The Red Junglefowl are elaborately coloured, with feathers ranging from dark red to light orange, whereas the White Leghorn birds are fixed for the dominant white locus and are pure white in colour. The intercross birds display an enormous range of plumage colouration, whilst genetically the advanced intercross gives far smaller confidence intervals for detected QTL than standard F_2_ intercrosses(DARVASI AND SOLLER 1995). The study presented here utilizes multiple generations for this analysis. A large scale QTL scan was performed using the F_8_ generation, whilst further targeted expression QTL (eQTL) studies were performed using the F_10_ and F_12_ generations to assess candidate genes identified in the QTL scan. By using a combination of targeted genetical genomics (whole genome transcriptomics of targeted individuals) to simultaneously map eQTL and correlate gene expression with intensity of red colouration, we identify 5 putatively causal genes affecting quantitative variation in red plumage colouration in the chicken.

## MATERIALS AND METHODS

### Study animals

The animals used in this study come from three separate generations (F_8_, F_10_ and F_12_) of an intercross between a Red Junglefowl male and three females from a White leghorn selection line. Food and water was supplied ad libitum, see (JOHNSSON *et al.* 2012) for further details of housing and the intercross. Each generation typically consists of around 100 individuals, with the exception of the F8 mapping population with 572 individuals reared in 6 batches. The F_10_ and F_12_ generations were also of approximately 100 individuals each, with 12 F_10_ and 12 F_12_ individuals used in this study.

### Colour phenotyping

Wings were removed post-mortem from individuals after slaughter at 212 days in the case of the F_8_ individuals. Each wing was photographed using a tripod mounted NIKON D5500 with a 40mm lens using a LF-PB-3 (Falcon Eyes) light tent and two fixed 24Watt DULUX L (Osram) lights mounted to brackets to ensure identical lighting conditions were used for each wing. A standardized blue background was used for each shot, whilst an X-rite colour checker chart (with both pure white and red coloured squares) was included in each photograph to normalize colour levels. Wing photos were taken on the blue background, and a corresponding image was taken of the blank background without the wing present. Raw image quality was used for subsequent calculations. Two separate phenotypes were extracted from each wing, peak and overall red intensity. Peak intensity was measured by selecting an area of approximately 1cm^2^ (∼14000 pixels) in the area with the strongest representative red intensity observed by eye in Adobe Photoshop CS6 and recording the median value from the red RGB channel. This value was then divided by the value measured from the ‘true’ red on the colour chart, to give a normalized peak red intensity for each wing. It is worth noting that since white is composed of full saturation in all channels this red value is negatively correlated with red colour, ie the darker red the feather is the lower the relative colour score will be and vice versa. The second phenotype uses the average red intensity measured across each wing. This was performed using the following steps:

### 1. Colour correction

The raw sensor measurements (pixel values without any of the usual processing of the camera) were extracted from the image file using the program dcraw (see http://www.cybercom.net/∼dcoffin/dcraw/ for the original version of the program and http://www.guillermoluijk.com/tutorial/dcraw/index_en.htm for a tutorial describing the program). For each of the images the location of the white patch of the colour checker with the highest intensity was automatically detected and the mean RGB vector over all pixels in this patch was computed. The colour vectors of all pixels in the image were then scaled (and converted to 16-bit unsigned integer) such that the resulting pixels in the white patch had all the same gray values. After this step all colour variations due to varying illumination and camera conditions were eliminated and all images could be compared.

### 2. Geometric correction and compilation of the wing mask

After normalization, colour correction registration points (essentially two crosshairs on the left and the right side of the blue background) were automatically detected in both images of an image pair. The locations of these registration points were then used to compute the geometric transformation that aligned the images without the wing with the image with the wing. Pixelpairs with blue colour in the background image and non-blue colour in the wing image define a mask indicating the location of the wing.

### 3. Extraction of red pixels

The mask computed in the previous step is used to set all non-wing pixels to a black background colour. Next the colour vectors in the masked image are converted from the RGB colour space to the HSV colour space using the Matlab function rgb2hsv. In this HSV the colour of a pixel is described by three values: H is the hue (described by an angle), S is the saturation described by an interval and finally V is the value, which is a non-negative number describing the lightness. In the application only reddish pixels are of interest. The corresponding region in colour space was defined as the region where the hue value was between 0.6 and 0.7 and the saturation value was greater than 0.75. Both the hue and the saturation values are located in the [0,1] interval. This procedure thus isolates all the red patterned feathers from the rest of the wing. After the pattern isolation, the red channel of the entire resulting trimmed image was measured in Photoshop. Therefore this second phenotype takes the median red intensity of every red area of the wing into consideration. The average value for this second phenotype is 184.4 with a standard deviation of 42.4 compared to the average value of 0.71 with a standard deviation of 0.18 for the single feather analysis used for the eQTL analysis (see below).

A separate eQTL fine mapping population was used for expression analyzes of budding feather tissue for candidate gene assessment in the previously detected QTL regions identified in the F_8_ generation. These birds were individuals from both the F_10_ and F_12_ generations from the advanced intercross (n=12 F_10_ and 12 F_12_ birds). Multiple generations were required to obtain a broad spectrum of red colour phenotypes. For these birds, growing feather buds were selected from adult wings for use in both phenotyping and gene expression assays. The feathers from each growing bud were removed and used in the phenotyping assay, to ensure that the gene expression and colour phenotype would match as closely as possible. These feathers were removed from the bud and stacked in 3 layers to mimic the wing (where feathers are overlapping). Red colour intensity was then measured using the same technique as for the F_8_ birds, with only the peak intensity measured. The developing basal root of the feather follicle (calami) was then used for RNA extraction and microarray analysis for eventual eQTL analysis.

### RNA extraction

5-6 growing feathers of 2-3 cm length were collected from each individual. The calamus was immediately cut from each feather and snap frozen in liquid nitrogen for RNA extraction, the rest of the feather was kept at −30 and used for colour measurements. RNA was extracted from a pooled sample of three to four calami per individual which had been homogenized in Lysing matrix D tubes(MP Biomedicals) in a FastPrep(MP Biomedicals) using TRIzol (Sigma Aldrich) reagent and following the manufacturers instructions. After quality checking the RNA with RNA Nano Chips(Agilent Technologies) in an Agilent 2100 Bioanalyzer to ensure that RNA was not degraded, the RNA was treated with DNAse I (Thermo scientific) followed by cDNA synthesis using a RevertAID KIT from Life technologies, and following the instructions provided with the kit. The cDNA was again quality checked using a Bioanalyzer 2100 and RNA Nano chips(Agilent Technologies) and subsequently labeled with a NimbleGen One colour labeling kit(Roche) according to the manufacturers instructions. Finally the cDNA was hybridized to NimbleGen 12×135k custom gene expression microarrays (Roche) and scanned using a NimbleGen MS200 Microarray scanner (Roche). The custom Microarrays have been used in previous publications by the group and contain all known Ensembl and RefSeq genes as well as EST probes (for microarray details see Johnsson et al. 2016). Microarrays were first preprocessed using DEVA software to normalize the data and all arrays were processed together.

### Genotyping, QTL mapping

DNA preparation was performed by Agowa GmbH (Berlin, Germany), using standard salt extraction. A total of 652 SNP markers were genotyped using an Illumina GoldenGate system, and gave a map of length ∼92675cM, with an average marker spacing of ∼ 16cM. QTL analysis was performed in R (R DEVELOPMENT CORE TEAM 2008) using the R/Qtl software package (BROMAN *et al.* 2003), with standard interval mapping and epistatic analyzes performed. Interval mapping was performed using additive and additive+dominance models. In the colour phenotyping QTL analysis batch and sex were always included as fixed effects. To control for potential family substructure, a Principal Component Analysis (PCA) of the genotype data was performed, and the first ten PCs fitted as covariates in the model, with all significant PCs retained in the final model (see (JOHNSSON *et al.* 2016) for further details). A sex interaction was also added, where significant. Digenic epistatic analysis was performed as per the guidelines provided in (BROMAN AND SEN 2009). Initially a global model was used that incorporated standard main effects and sex interactions. Epistasis was then built on this model, starting with the most significant loci and working down for each trait. Significance thresholds were calculated by permutation (CHURCHILL AND DOERGE 1994; DOERGE AND CHURCHILL 1996), with thresholds of 20% and 5% genome wide significance being the cut-offs for suggestive and significant loci respectively. This corresponded to approximately LOD cut offs of 3.6 and 4.4. Confidence intervals were calculated using a 1.8 LOD drop method (MANICHAIKUL *et al.* 2006). Epistasis thresholds were calculated in a similar manner, with 20% and 5% genome-wide thresholds used.

### eQTL Mapping

The individuals in the F_10_ and F_12_ fine mapping eQTL population were genotyped for sixteen SNP markers which had been identified in the F8 mapping population as being flanking markers for the QTL with the two colour phenotypes (see supplementary table 1 for locations and primers). Five were located on chr2, three on chr10 and chr14 respectively, and two markers on chr 15 and chr 11 and finally one marker on chr26, for a total of 16 markers. These SNP markers were all genotyped by pyrosequencing on a QIAGEN pyromark q24 using QIAGEN GOLD reagents. DNA was extracted from blood using a standard salt extraction protocol, except for a small number of individuals were blood samples were missing and DNA was instead extracted from feathers using a modified protocol for the Qiagen DNEasy kit with dithiothreitol(DTT) added to the lysis step.

Analysis was performed using a three-step procedure. Initially, all genes that were present in the colour QTL regions detected in the F_8_ analysis were selected as candidates (n=875, Reference genome version GalGal4). These probes were then correlated with the specific red colour phenotype obtained for each feather sample (see above), using a linear model to fit gene expression levels and colour score using the lmFit function in the limma R package (RITCHIE *et al.* 2015; R STUDIO TEAM 2016). All probes that had a significant association between gene expression and colour score at a FDR significance level of 0.05 were retained (n= 76, see supplementary table 2). The last step was to perform eQTL mapping in R/qtl, using standard interval mapping, with sex and batch (F_10_ or F_12_) included as fixed factors. All the genes from stage 2 were tested against the regions genotyped using the 16 SNP markers mentioned above (on chromosomes 2, 10, 11, 14, 15 and 26). Permutation testing was used to set an experiment-wide threshold for each gene (1000 permutations per gene), with a 5% significance and a 20% suggestive threshold obtained (LOD values of ∼3.0 and ∼2.2 respectively). To control for the total number of genes (n=76), a principal component analysis was first performed on these 76 genes. 7 significant eigenvalues accounted for 97% of the total variation in the data. Thus, as it is only required to control for the number of independent tests, a multiple testing correction of 7 was added to each LOD threshold. Using a log_10_ transformation to convert this to a LOD score, this gives an increase of 0.85 LOD to each threshold, meaning an eQTL was suggestive with a LOD score of ∼3.05, and significant with a LOD score of ∼3.85.

## RESULTS

### QTL analysis

QTL analyzes were performed using both peak red intensity and overall red intensity in the F_8_ birds (n= 380). Three separate QTL were identified for peak intensity on chromosomes 2 at 149cM, chromosome 10 at 176cM, and chromosome 14 at 207cM. In the case of the loci on chromosomes 2 and 10, the RJF genotype led to an increase in red intensity, whilst for the chromosome 14 locus the WL genotype led to an increase in red intensity. Four separate QTL were identified for overall red intensity on chromosomes 2 (at 81cM), 11 (at 73cM), 15 (at 148cM) and 26 (at 0cM). Once again, for two of the loci (on chromosomes 2 and 15) the RJF genotype led to an increase in red intensity. The other loci on chromosomes 11 and 26 mainly acted through dominance rather than additive variation.

### Targeted Expression Analyses

The loci that were identified using the initial genome-wide F_8_ scan were then assessed using gene expression data from the F_10_ and F_12_ generations. We correlated the expression levels to the relative intensity of red colouration with all genes (a total of 875 probesets) located within the confidence intervals of the seven QTL regions. After multiple testing corrections (FDR, p=0.05) were applied we found 76 probesets that were correlated with the peak intensity of red colouration. (FDR adjusted p-value < 0.05).

In order to further narrow down the list of potential candidate genes we performed an eQTL analysis using the probesets that correlated with colour using a set of targeted markers within colour QTL. We detected six eQTLs covering 5 genes that were genome-wide suggestive (including a multiple testing correction for the total number of genes tested), two for genes associated with overall red intensity and four that are associated with the peak red intensity (see table 2). All were trans-eQTLs, being controlled from a region on chromosome 10 (15Mb).

**Table 1.**
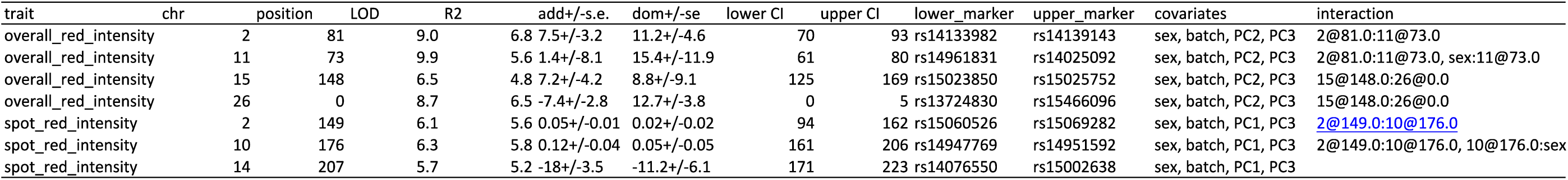
Results from the QTL scan of the two phenotypes(red spot intensity and average red red) in the initial F8 mapping population. QTL locations (in cM), Lod scores, effect sizes (r^2^), additive and dominance values, confidence intervals (as per a 1.8 lod drop method), covariates and interactions are all shown.

**Table 2.** Summary of candidate genes with both a significant correlation with red intensity and a significant eQTL. The table includes the log of Fold change for the probe in question, t value of the gene expression, the adjusted p-value and the B value for the correlation between expression levels and colour score. In addition, it has the chromosomal location of the eQTL marker (plus the marker name), the additive and dominance effects of the eQTL, the % of variation explained by the eQTL (r-squared value), lod score and the location of the gene itself.

| colour correlation |  |  |  |  | eQTL |  |  |  |  |  |  |
| --- | --- | --- | --- | --- | --- | --- | --- | --- | --- | --- | --- |
| Gene | logFC | t | adj.P.Val | B | eQTL origin | eQTL marker snp | additive effect (+/- SE) | dominance effect (+/- SE) | % of variation | lod | Gene location |
| WDR24 | 4.52 | 3.20 | 0.05 | -1.83 | Chr 10 | Gg_rs14949856 | 0.9190 (0.3162) | 1.9981 (0.4790) | 51.12 | 3.73 | 14 |
| PHLDA3 | 4.04 | 3.12 | 0.05 | -1.97 | Chr 10 | Gg_rs14949856 | 0.8264 (0.2938) | 1.7414 (0.4451) | 48.37 | 3.45 | 26 |
| LAD1 (ensgal31450) | 2.87 | 2.85 | 0.05 | -2.51 | Chr 10 | Gg_rs14949856 | 0.6549 (0.2196) | 1.2526 (0.3327) | 48.06 | 3.41 | 26 |
| CREBBP | 4.24 | 3.06 | 0.05 | -2.13 | Chr 10 | Gg_rs14949856 | 0.8353 (0.3221) | 1.8305 (0.4879) | 45.71 | 3.18 | 14 |
| ARL8A | 3.04 | 2.99 | 0.05 | -2.23 | Chr 10 | Gg_rs14949856 | 0.5042 (0.2297) | 1.3767 (0.3480) | 45.82 | 3.19 | 26 |
| LAD1 (ensgal0410) | 4.39 | 2.90 | 0.05 | -2.40 | Chr 10 | Gg_rs14949856 | 0.9748 (0.3519) | 1.8517 (0.5331) | 44.18 | 3.04 | 26 |

### A network of trans-eQTL between colour QTL

We see a trans regulation of candidate gene expression, with a locus on chromosome 10 controlling multiple target sites. The most significant trans-eQTL effect originates from chromosome 10 and controls two genes on chromosome 14 *(CREBBP* and *WDR24*), and three genes on chromosome 26 (*ARL8A, PHLDA3,* and *LAD1*), see figure 2. There is also a selective sweep (RUBIN *et al.* 2010) located within the chromosome 10 QTL region around 500kb downstream from our eQTL marker location and within the confidence interval for the colour QTL identified in the F_8_ population.

**Figure 1.**
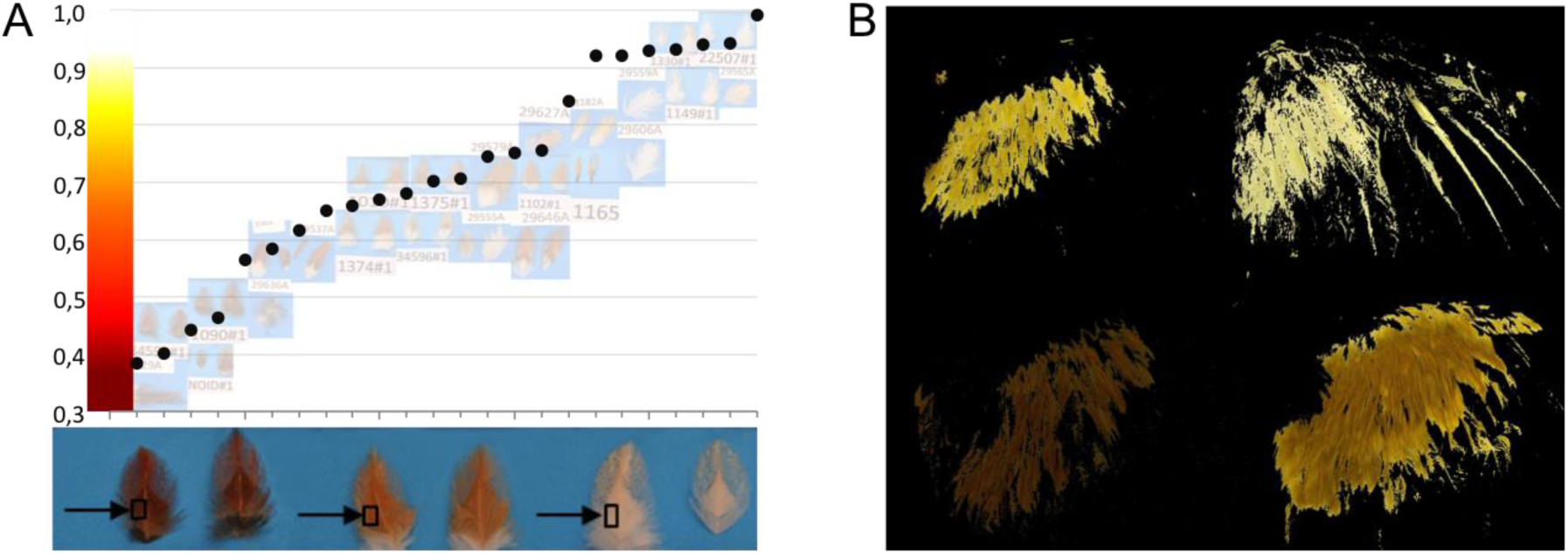
Quantitative colour phenotypes used for QTL/eQTL mapping. A) Peak red intensity. The feathers that we have used in the gene expression measurements are placed on a blue background and their level of red intensity is measured relative to a True red colour chart. The darkest red feathers score between ∼0.4-0.5 and the almost pure white feathers score between ∼0.9-0.99. B) Depicts four representative measurements of Average red colour score measured across the right wing.

**Figure 2.**
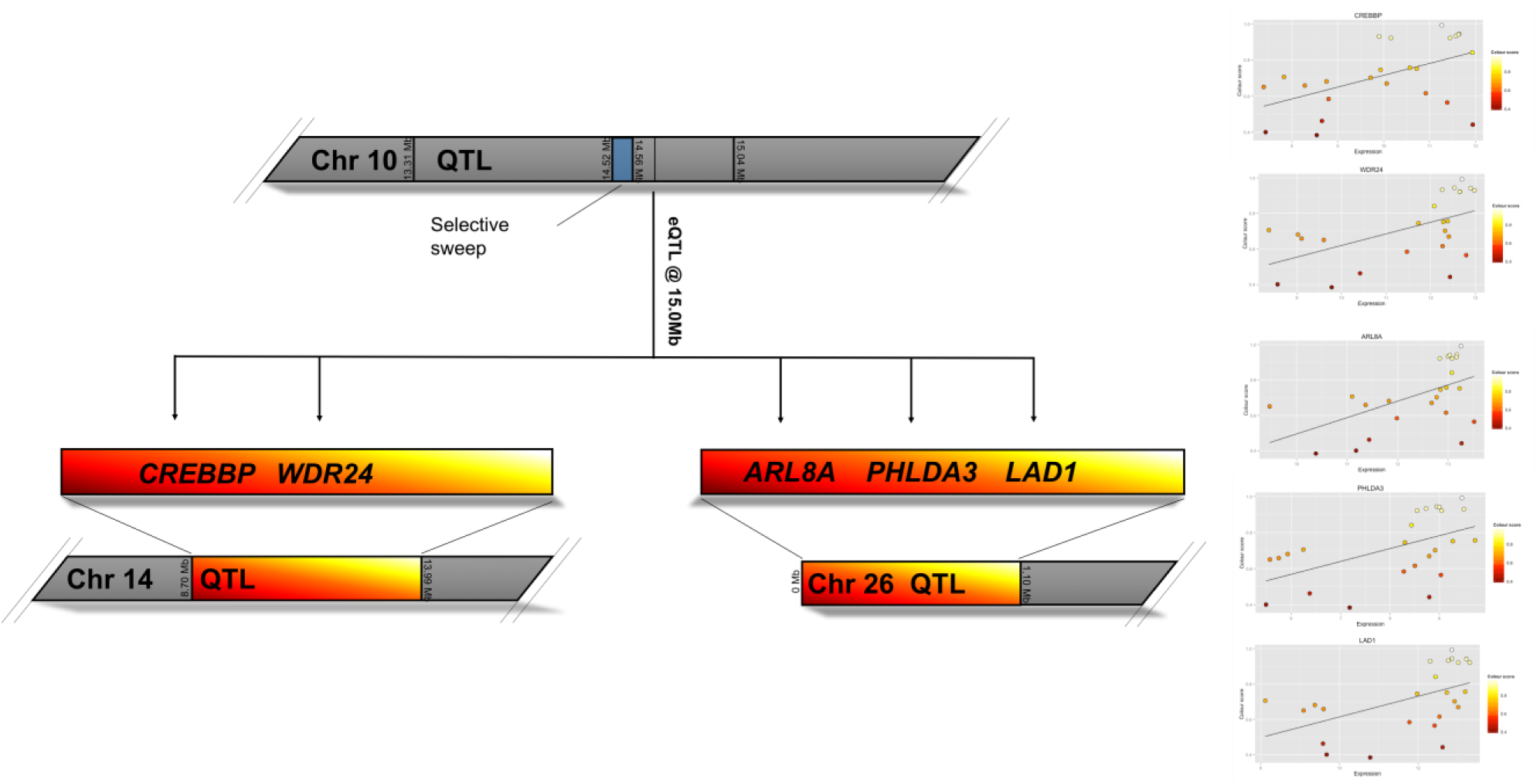
Cartoon depicting the genetic regulation of gene expression for the candidate genes with both a correlation with red intensity and a significant eQTL.

## DISCUSSION

The aim of this study was to identify genes and genetic loci that affect red colouration in the chicken. By treating colour as a quantitative trait, as opposed to looking for solely discrete classes, we have identified a total of seven small effect QTL that until now have been overlooked. Furthermore, by combining this with a targeted genetical genomics approach in an additional population we have identified five candidate genes affecting the intensity of red colouration. These genes would be unlikely to have been identified using more classical (presence/ absence of major genes) linkage methods. This is unsurprising given that the average effect size was around 7% in our study, whilst the high degree of epitasis between loci would have meant they would have been extremely hard to pinpoint using single scan approaches. In particular chromosome 10 appears to have a strong effect, controlling loci on chromosomes 14 and 26. It therefore appears that this locus has trans acting effects in combination with epistasis. This locus also contains a selective sweep that is fixed in domestic layer birds (the majority of which are white), yet segregating in wild RJF. The presence of this selective sweep adds weight to the evidence that this locus truly affects colour intensity.

Several genes have been identified as affecting colour in the chicken, however these are all major-effect genes that have been identified using a classical linkage/ single gene approach. In the chicken, the Dominant White locus (on chromosome 33, an allele of the *PMEL* gene) (KERJE *et al.* 2004), the classic extended phenotype (E) linked to *MC1R* on chromosome 11 at 19Mb (TAKEUCHI *et al.* 1996; KERJE *et al.* 2003), and the Dark Brown (Db) locus on chromosome 1 (linked with SOX10) (GUNNARSSON *et al.* 2007) all affect pigmentation (white/black in the case of Dominant White and E, and red in the case of Dark brown), whilst sex-linked barring (on chromosome Z) and the *EDNRB2* (panda) locus in quails (MIWA *et al.* 2007) primarily affect patterning. Despite these examples, studies of pigmentation inheritance in crosses imply that many more genes have yet to be identified. Specifically, to examine buff, brown and red colouration, crosses have been made with a variety of breeds, including Junglefowl x Buff Minorca and Rhode Island Red breeds (BRUMBAUGH AND HOLLANDER 1966), Brown and Light Brown Leghorn breeds (SMYTH JR 1965), Dark Cornish and Black breasted Red (KIMBALL 1955), and Villafranquina birds (CAMPO AND ALVAREZ 1988). These studies revealing a complex pattern of inheritance and multiple modifier loci exist that affect pheomelanine. For example with the buff phenotype, four autosomal factors were proposed, named ginger, mahogany, dilute and champagne blond (BRUMBAUGH AND HOLLANDER 1966), whilst many studies noted many variations in the intensity of the red colouration present, even when segregating for a known major effect gene (SMYTH JR 1965) (CAMPO AND ALVAREZ 1988). The wide variety of phenotypes that rapidly emerge indicate that multiple epistatic interactions and numerous genes of small effect is the norm, yet such modifier loci have yet to be identified.

The QTL that we report here overlap none of these previously identified major effect loci. The genes and QTL are all novel loci (as affecting colour), though in the case of the QTL on chromosome 11, this is around 4Mb downstream of the *MITF* gene. It is possible that this locus is a modest trans acting factor affecting the *MITF* gene, however no correlation was found between any of the colour measurements and *MITF* expression. Interestingly, the linkage between this locus and *MITF* may have even caused the effects to be erroneously ascribed to *MITF* alleles. When considering the classical studies it is perhaps logical that these modifier loci are discrete to the genes identified previously. For example Dominant White is known to be ineffective against red pheomelanin, whilst the silver allele for sex-linked silver (PMEL) has no discernible modifying effect on red in females (KIMBALL 1955). Our study indicates that by concentrating solely on such major genes, which are easier to identify, much variation is missed.

Of the candidate genes identified, particular interest comes from the interaction and correlation seen between the gene *CREBBP* and red intensity. *CREBBP* is controlled by the trans-acting eQTL locus on chromosome 10 that also contains the selective sweep. *CREBBP* binds to *CREB*, whilst *CREB* itself is a transcription factor strongly involved in melanin synthesis, principally by binding and activating *MITF* via the cAMP response element (SAHA *et al.* 2006). Given this role of *CREB* and its close interplay with *CREBP*, it therefore represents an excellent candidate as small-effect modifier of red colouration via melanogenesis to alter pheomelanine. Currently, *CREBBP* has primarily been found to be involved with neuronal growth and development (SHARMA *et al.* 2010; LOPEZ-ATALAYA *et al.* 2011), so a role in feather colour expression is relatively novel. Of the other candidate genes identified, *WDR24* is a master regulator of lysosome function. Given one function of lysosomes is to regulate and degrade pigment molecules, this also represents another excellent candidate gene. Similarly, the gene *ARL8A,* an acetylated Arf-like GTPase, is localized to lysosomes and affects their mobility (HOFMANN AND MUNRO 2006). Of the remaining genes, *PHLDA3* is implicated in tumour suppressor function, is a target of p53 and also impedes somatic cell reprogramming (QIAO *et al.* 2017), whilst *LAD1* encodes ladinin-1, a collagenous anchoring filament protein that is involved with the maintenance of cohesion at the dermal-epidermal boundary (TEIXEIRA *et al.* 2015), and defects in this gene are related to linear IgA disease, an autoimmune blistering disease (ISHIKO *et al.* 1996).

In summary, by treating the intensity of red colouration as a purely quantitative trait and utilising a variety of advanced intercross generations and targeted gene expression profiling we are able to identify multiple small-effect loci that interact to determine the red colouration present in RJF. Using a targeted genetical genomics approach we are able to identify several high quality candidate genes that appear to have direct relevance to the melanogenesis pathway and can thus modify overall red intensity. As far as we are aware this is the first study to identify QTL and candidate genes that can more subtly modify the intensity of red colouration in feathers, with such small effect loci being potentially far more prevalent in wild populations.

## ACKNOWLEDGEMENTS

The research was carried out within the framework of the Swedish Centre of Excellence in Animal Welfare Science, and the Linköping University Neuro-network. SNP genotyping was performed by the Uppsala Sequencing Center. The project was supported by grants from the Carl Tryggers Stiftelse, Swedish Research Council (VR), the Swedish Research Council for Environment, Agricultural Sciences and Spatial Planning (FORMAS), and European Research Council (advanced research grant GENEWELL 322206).

## FIGURE LEGENDS

**Supplementary table 1.**
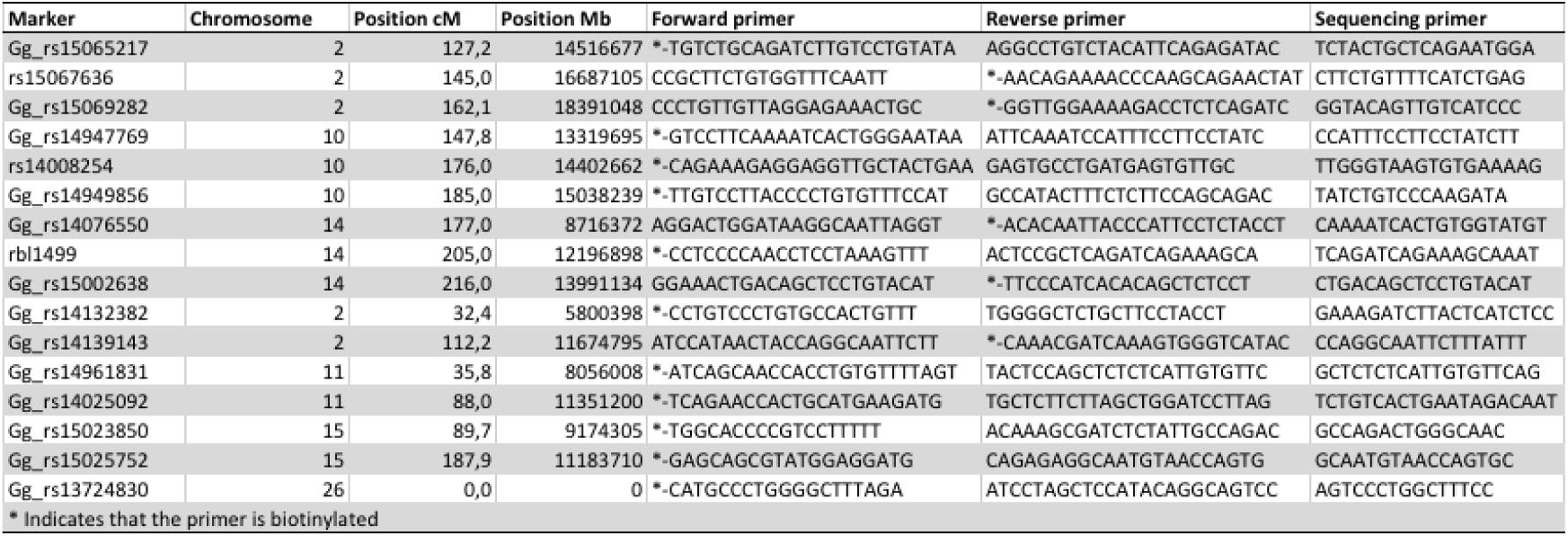
The table contains relevant information about the SNP markers and their respective primers used in the eQTL analysis. Position is both given in cM which relates to the QTL intevals, the Mb position is from the chicken reference genome(GALGAL4) and the position is given for the focal SNP.

**Supplementary Table 2.**
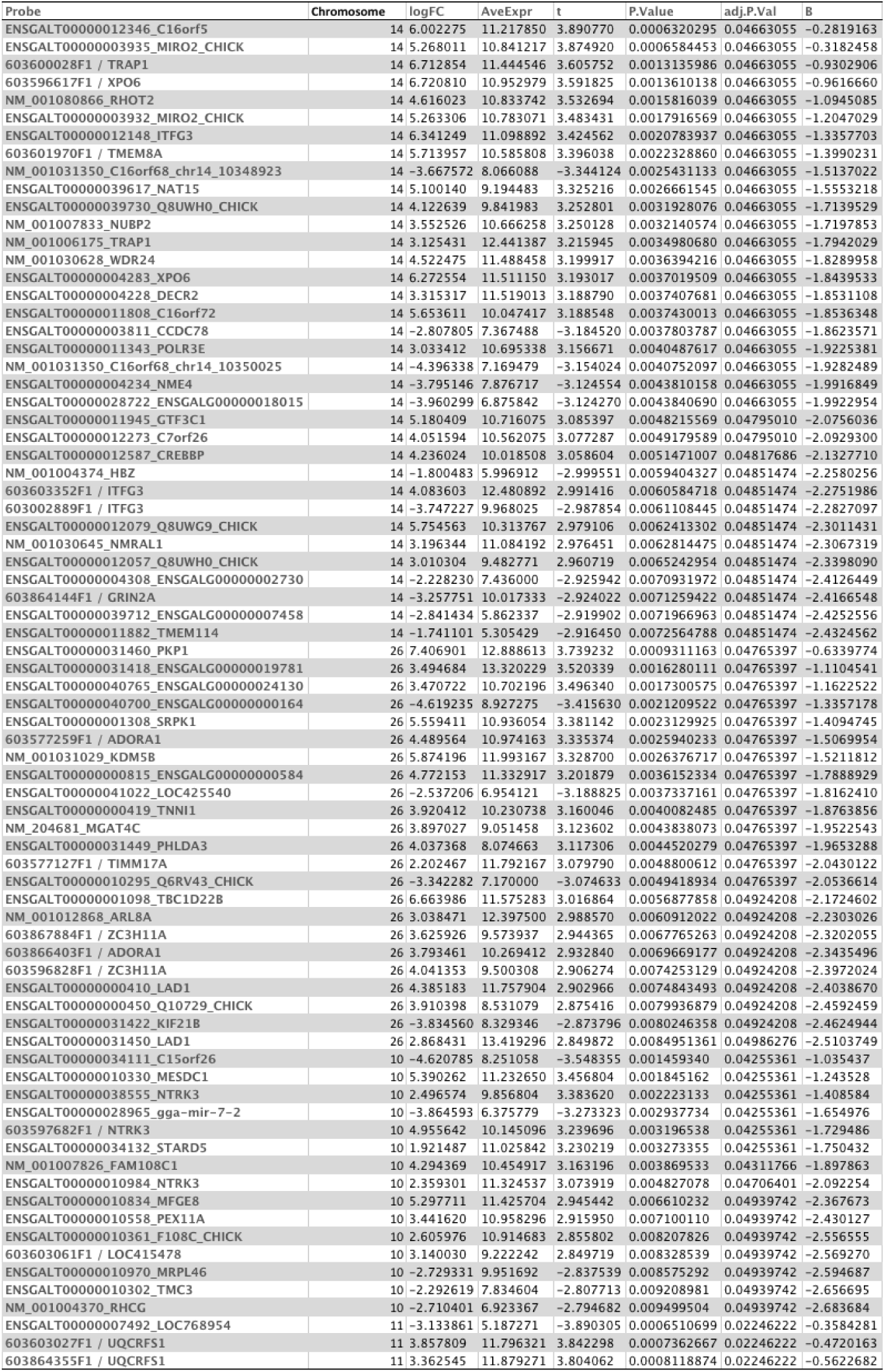
Differentially expressed genes that correlate with the intensity of red colouration and have an FDR adjusted p-values < 0.05. The table also contains information about Log fold change as well as average expression for each probe.

